## Supplementary figures and images for "A chromosome-level reference genome for Pacific herring (*Clupea pallasii)* from the Bering Sea"

### Supplemental Figure 1

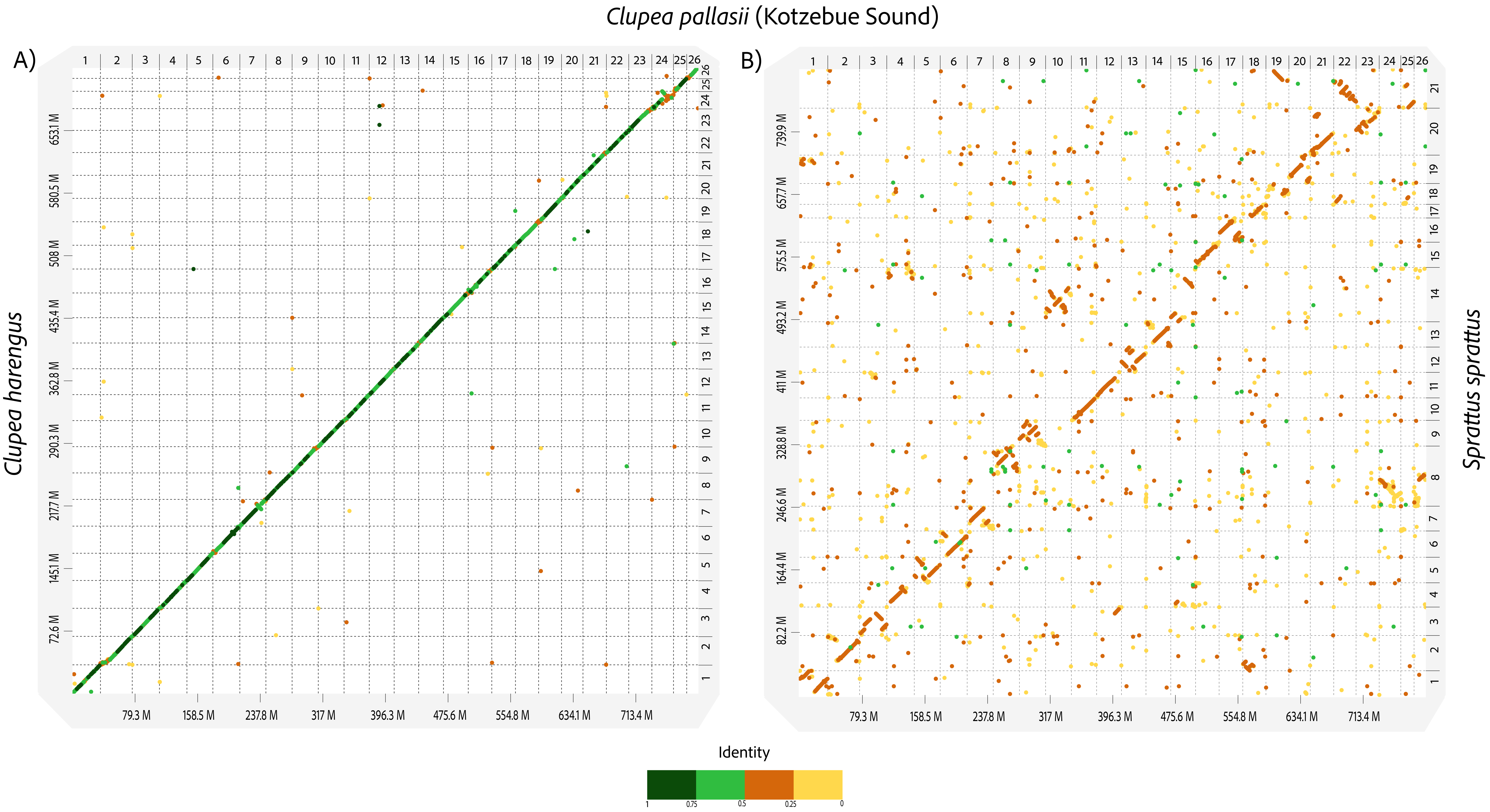

### Supplemental Figure 2

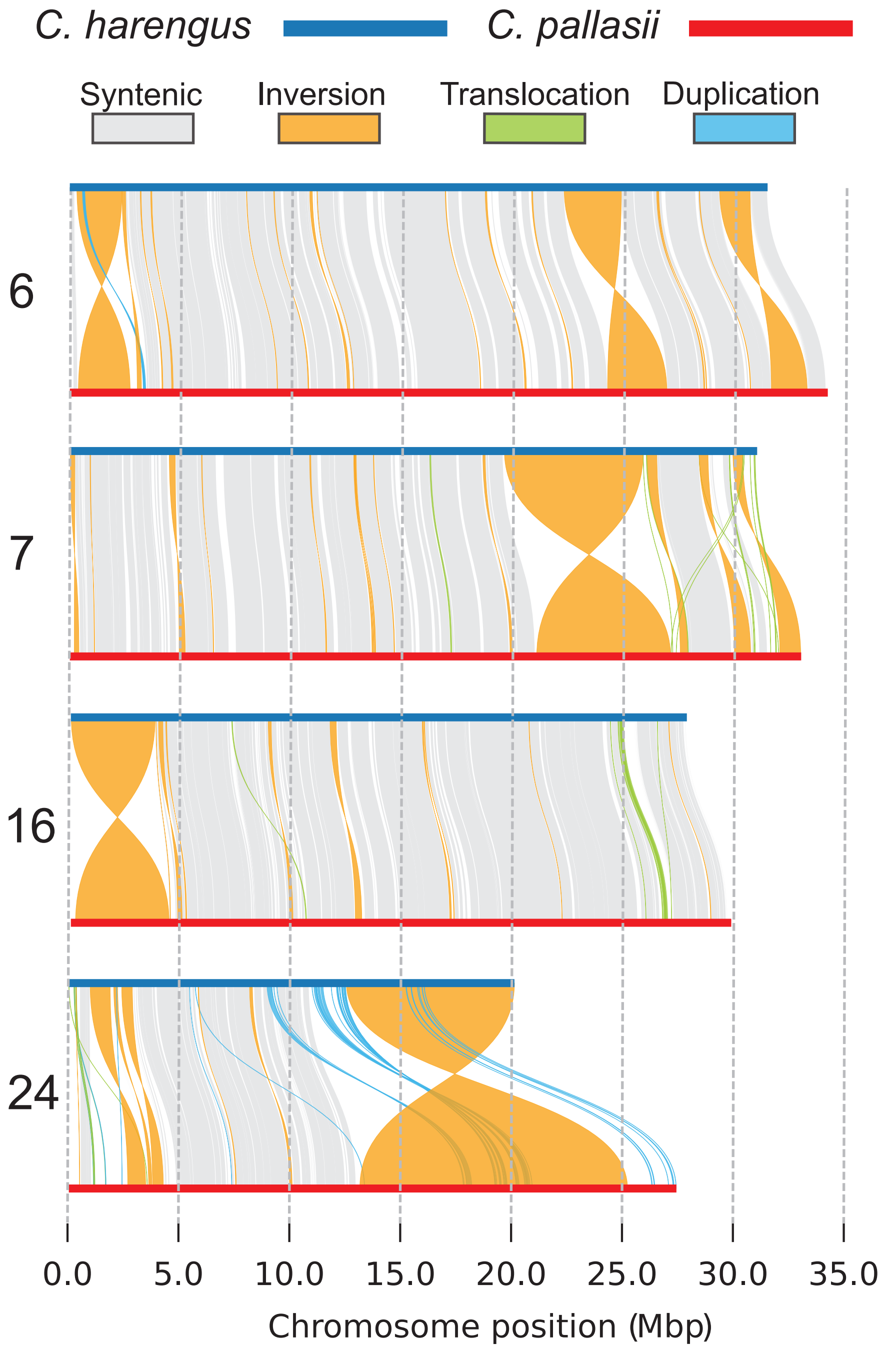
